# Recombining without Hotspots: A Comprehensive Evolutionary Portrait of Recombination in Two Closely Related Species of Drosophila

**DOI:** 10.1101/016972

**Authors:** Caiti S. Smukowski Heil, Chris Ellison, Matthew Dubin, Mohamed A. F. Noor

## Abstract

Meiotic recombination rate varies across the genome within and between individuals, populations, and species in virtually all taxa studied. In almost every species, this variation takes the form of discrete recombination hotspots, determined in Metazoans by a protein called PRDM9. Hotspots and their determinants have a profound effect on the genomic landscape, and share certain features that extend across the tree of life. Drosophila, in contrast, are anomalous in their absence of hotspots, PRDM9, and other species-specific differences in the determination of recombination. To better understand the evolution of meiosis and general patterns of recombination across diverse taxa, we present what may be the most comprehensive portrait of recombination to date, combining contemporary recombination estimates from each of two sister species along with historic estimates of recombination using linkage-disequilibrium-based approaches derived from sequence data from both species. *Using Drosophila pseudoobscura* and *Drosophila miranda* as a model system, we compare recombination rate between species at multiple scales, and we replicate the pattern seen in human-chimpanzee that recombination rate is conserved at broad scales and more divergent at finer scales. We also find evidence of a species-wide recombination modifier, resulting in both a present and historic genome wide elevation of recombination rates in *D. miranda*, and identify broad scale effects on recombination from the presence of an inter-species inversion. Finally, we reveal an unprecedented view of the distribution of recombination in *D. pseudoobscura*, illustrating patterns of linked selection and where recombination is taking place. Overall, by combining these estimation approaches, we highlight key similarities and differences in recombination between Drosophila and other organisms.

**Author Summary:** Recombination, or crossing over, describes an essential exchange of genetic material that occurs during egg and sperm development and has consequences for the proper segregation of chromosomes, and for the evolution of genomes and genomic features. In our study, we compare genome wide recombination rate in two closely related species of the fruit fly Drosophila to understand if and how recombination changes over time. We find that recombination does indeed change, we observe globally increased recombination in one species, and differences in regional recombination likely reflecting the result of a chromosomal rearrangement in both species. Moreover, we show that the extent that recombination changes is dependent on the physical scale at which recombination is measured, likely reflecting selection pressures on recombination distribution and replicating a pattern seen in human-chimpanzee recombination. Apart from between-species differences, we note several ways in which the Drosophila recombination landscape has changed since Drosophila diverged from other organisms. In contrast to species of fungi, plants, and animals, Drosophila recombination is not concentrated in discrete regions known as hotspots, nor is it increased near the start of genes, suggesting that despite the importance of the recombination process, the determinants of recombination have been shifting over evolutionary time.

## Introduction

Homologous meiotic recombination is a crucial mechanistic and influential evolutionary process. The creation of a physical link between homologous chromosomes during meiosis I promotes proper segregation of chromosomes, preventing aneuploidy and cell death. Evolutionarily, recombination breaks down linkage between loci, allowing sites to segregate independently, and thereby facilitating adaptation and the purging of deleterious mutations. Indeed, recombination is often touted as the fundamental benefit of sexual reproduction. Nonetheless, measuring recombination elicits many challenges. Researchers have used a myriad of ways to estimate recombination rate, which can broadly be grouped into two categories: assessing direct, current rates or capturing indirect, historical rates (1, 2). The most common approach for assessing direct, current rates entails genotyping individuals from known pedigrees or controlled crosses, which provides a coarse estimate of recombination rate in a single meiosis or small number of meioses. However, this method is costly, laborious, and very difficult to achieve fine scale resolution, often involving a trade-off between number of markers and number of individuals to survey. In contrast, with the rise of population sequencing data over the past decade, a statistical approach has been developed and now widely used that estimates the population recombination parameter, ρ, to computationally infer genome wide, population-averaged recombination rate (3-5). The approach can achieve very fine scale resolution, unmasking genomic features and leading to the discovery of determinants of recombination rate variation.

The development of recombination maps in a wide range of species through both direct and indirect methods has dramatically enhanced our understanding of where recombination takes place in the genome, what factors are responsible, and the selective pressures governing changes in recombination rate over time. For example, one of the most fundamental patterns observed is the existence of variation in recombination rate across the genome, and between populations and species in every organism examined (6-9). Moreover, this variation in recombination rate is non-random, and dependent upon the physical scale at which it is examined. For instance, in many organisms, the broad scale recombination landscape is shaped by large scale structural features of the chromosomes, while the fine scale variation manifests as “hotspots”: discrete genomic regions, typically 1-2 kilobases (kb) in length in which recombination increases a magnitude fold above the background rate (2, 10-12).

With an increasing number of recombination maps developed for related species, it has become apparent that selective pressures governing recombination rate are also dependent on physical scale. Recombination rates observed at the megabase scale or greater exhibit conservation between closely related species (6, 8, 13-18), while recombination hotspots show nearly no overlap between closely related species (19-21). Some of the conservation at broad scales may be due to the necessity for at least one crossover per chromosome to ensure proper segregation of chromosomes during meiosis (8, 22-24). This mandates a minimum level of recombination, as a crossover rate that is too low is likely to be highly deleterious. In contrast, too much recombination breaks apart advantageous allele combinations, threatens genomic integrity, and may introduce mutations at hotspots (2, 12, 25). Therefore, on a broad scale, recombination is likely constrained between a lower bound dictated by chromosome segregation, and an upper bound determined by maintaining genomic integrity. At fine scales, the recombination landscape is shaped by a phenomenon known as the “hotspot paradox” (26-28), which describes the self-destructive nature of the resolution of meiotic crossover products. A method to counteract the loss of hotspots was unknown until the recent discovery of the zinc finger histone-methyltransferase, *Prdm9*, which controls the distribution of recombination in mice and humans (29, 30). Positive selection on *Prdm9*’s zinc fingers has created new DNA binding residues, thereby initiating new populations of hotspots (29, 31-36), and resulting in rapid turnover of hotspots and fine scale recombination rate divergence between individuals, populations, and species who have *Prdm9* (16, 21, 31-34, 36-41).

Furthermore, *Prdm9* may also determine the distribution of recombination events across the genome, possibly directing recombination machinery away from promoters and other functional genomic elements (33, 40, 42). In some of the organisms lacking *Prdm9* such as dogs, plants, and yeast, (35, 43, 44), hotspots are often found overlapping promoter regions/transcription start sites (TSS), presumably due to the open chromatin structure allowing recombination machinery access to the DNA (11, 45-50). These observations set up a model in which organisms without *Prdm9* are opportunistic: they form recombination hotspots in “windows of opportunity,” most often nucleosome depleted regions around promoters/TSS. On the other hand, in organisms with *Prdm9*, *Prdm9* most likely functions to introduce meiotic specific H3K4me3 modifications and directs recombination machinery to initiate at these locations away from genomic elements (51, 52).

Drosophila stands out from these systems in its unique recombination landscape. Like yeast and plants, Drosophila are missing *Prdm9* (53), but unlike these species, Drosophila lack highly localized recombination hotspots, the only system known to be missing such hotspots besides *C. elegans* (54-59). Moreover, Drosophila recombination is sex-specific (males don’t experience crossing over), and chromosome pairing and synapsis proceed normally in its absence (60-62). Drosophila recombination poses an intriguing system to further explore such questions as: How is recombination distributed in a species lacking hotspots? Are recombination rates conserved or divergent at varying scales? Are there additional differences between Drosophila recombination and recombination in other taxa?

To address these questions, we have developed both empirical and LD-based estimates of recombination in each of two closely related species, *Drosophila pseudoobscura* and *Drosophila miranda,* to better understand the recombination landscape and how it changes over evolutionary time. Empirical estimates of recombination rate are available for both species from McGaugh et al. (63), and here, we present genome wide LD-based estimates of recombination (via LDhelmet (64)). We show that LD-based recombination maps generally resemble current recombination rates in these species, and demonstrate general conservation between species over three million years divergence. We find that *D. miranda* recombination rates have been and still are higher than those of *D. pseudoobscura*, suggestive of a global recombination modifier. We uncover potential historic effects of an inversion on broad-scale recombination rate, and similar to mice and humans, recombination rates are more conserved between species at larger genomic scales (>100kb) than finer genomic scales. Our fine scale recombination rate estimates illuminate where recombination takes place in the Drosophila genome, and confirm the lack of highly localized recombination hotspots in this taxon. In total, this work highlights key similarities and differences of Drosophila recombination compared to other organisms, adding a more thorough understanding to the evolution of meiosis and recombination across disparate taxa.

## Results and Discussion

### How well do Linkage Disequilibrium-based linkage maps reflect present-day measures of recombination?

The use of statistical programs on population sequencing data has drastically accelerated the study of recombination rate variation, particularly at fine scales. However, while Linkage Disequilibirum (LD)-based inference of recombination rate is a powerful tool, this method has its drawbacks too: linkage disequilibrium is influenced by many other factors apart from recombination, including demography, selection, and genetic drift (5). Additionally, linkage disequilibrium represents a historical population averaged recombination rate while empirical methodologies typically survey only one population for one generation of meiosis. Thus, LD-based maps can potentially deviate from direct, single-generation empirical estimates of recombination for several reasons (65-68). Therefore, to first clarify how well our LD-based maps replicate empirical recombination rates, we created genome wide LD-based recombination maps for *D. pseudoobscura* and *D. miranda*. The LD-based recombination maps were made with the program LDhelmet (64), similar to the widely used program LDhat (4), but explicitly designed for Drosophila to account for biological and technical differences in the datasets, such as a much higher background recombination rate than human. We confirmed that estimates of recombination from LDhelmet correlated strongly with a simple direct measure of linkage disequilibrium decay in our dataset (**S1 Fig.**). We then compared the LD-based recombination estimates to those from the empirical maps generated by McGaugh *et al*. (63). The empirical maps were generated by genotyping SNP markers at 150-200 kb intervals for several of the largest chromosome groups for two populations of *D. pseudoobscura*, and one of *D. miranda*, and several selected regions of 20kb and 5 kb of chromosome 2 for one population of *D. pseudoobscura* (55).

Recombination rates obtained from these two strategies are significantly correlated at a broad scale (˜200kb, **Table 1, Fig. 1, S2-S4 Figs**), although empirical-LD-based correlations are generally stronger in *D. pseudoobscura*. A possible explanation for this difference is that LD-based rates typically don’t capture extreme values seen in the empirical landscape (**Fig. 1**, **S2-S4 Figs**), which may explain why the correlation is weaker in the elevated recombination map of *D. miranda*. The particular strains used in the *D. miranda* empirical study may also have an atypical recombination distribution relative to the historical population, or that there could be greater among-strain variation across *D. miranda*. *D. miranda* XLgroup1a in particular stands as an outlier (r=0.227), although the empirical map in this region is broken into three pieces due to failure of markers, potentially introducing more error into these estimates. We also have empirical estimates for a small portion of *D. pseudoobscura* at finer scales of 20kb and 5kb; the correlation between empirical and LD-based maps persists at 20 kb (r=0.68, **S5 Fig.**), but there is too little data at the 5kb scale to be conclusive (**S6 Fig.**).

**Fig. 1.**
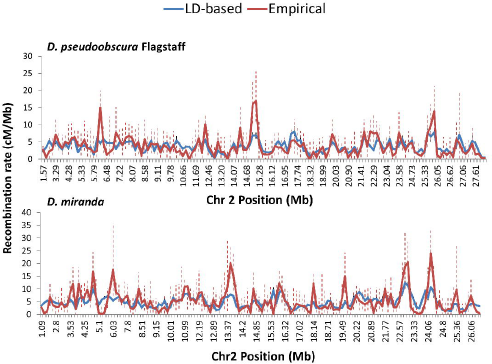
A comparison of empirical and LD-based estimations of recombination rate. Empirical recombination rates (red) are plotted with LDhelmet derived recombination rates (blue) across chromosome 2 for *D. pseudoobscura* Flagstaff (top) and *D. miranda* (bottom). All recombination rates are reported in cM/Mb. Error bars depict 95% Confidence Intervals for the empirical (red, dashed), and LD-based (black, solid).

**Table 1.** Empirical vs. LD-based comparisons of recombination rate. Empirical recombination rates were obtained from McGaugh *et al* 2012. LD-based recombination rates were averaged over empirical windows and converted to cM/Mb (see Methods). A dash indicates not enough data to assess the correlation.

|  | Interval size<br>Average,<br>Median<br>(Mb) | Number<br>of<br>intervals | Average<br>empirical<br>recombination<br>rate (cM/Mb) | Average LD-based<br>recombination<br>rate (cM/Mb) | Correlation<br>(Pearson's<br>r, p-value) |
| --- | --- | --- | --- | --- | --- |
| **2** |  |  |  |  |  |
| *D. pseudoobscura* Flagstaff | 0.215, 0.142 | 140 | 3.55 | 3.71 | 0.624,<br>p<0.0001 |
| *D. pseudoobscura* Pikes Peak | 0.194, 0.149 | 158 | 3.47 | 3.75 | 0.658,<br>p<0.0001 |
| *D. miranda* | 0.189, 0.147 | 154 | 4.85 | 4.87 | 0.639,<br>p<0.0001 |
| **XLgroup1a** |  |  |  |  |  |
| *D. pseudoobscura* Flagstaff | 0.215, 0.148 | 7 | 3.76 | 5.17 | - |
| *D. pseudoobscura* Pikes Peak | 0.211, 0.198 | 24 | 3.32 | 3.47 | 0.657,<br>p=0.0005 |
| *D. miranda* | 0.247, 0.202 | 18 | 4.95 | 5.82 | 0.227,<br>p=0.1806 |
| **XRgroup6** |  |  |  |  |  |
| *D. pseudoobscura* Flagstaff | 0.164, 0.143 | 68 | 3.94 | 3.66 | 0.710,<br>p<0.0001 |
| *D. pseudoobscura* Pikes Peak | 0.166, 0.146 | 71 | 4.44 | 4.55 | 0.719,<br>p<0.0001 |
| *D. miranda* | 0.163, 0.149 | 74 | 5.40 | 5.67 | 0.506,<br>p<0.0001 |
| **XRgroup8** |  |  |  |  |  |
| *D. pseudoobscura* Flagstaff | 0.269, 0.172 | 32 | 4.52 | 4.03 | 0.750,<br>p<0.0001 |
| *D. pseudoobscura* Pikes Peak | 0.248, 0.209 | 36 | 4.72 | 4.47 | 0.682,<br>p<0.0001 |
| *D. miranda* | 0.200, 0.187 | 34 | 6.12 | 6.06 | 0.403,<br>p=0.0182 |

Few studies have compared empirical and LD-based estimates, but similar results were obtained from LDhelmet in *D. melanogaster* (Raleigh, NC population; ˜200kb: 2L r=0.73, 2R r=0.75, 3L r=0.61, 3R r=0.73, X r=0.71 (64)). Chan *et al.* used data from Singh *et al.* (57) and FlyBase (69), at resolutions of approximately 100-200kb, so we present the first data obtained from LDhelmet demonstrating correspondence in LD-based and empirical rates at scales under 100kb. One of the only other comparisons is from human, where correlations between LD-based and empirical recombination rates are nearly replicates at the megabase scale (r=0.98)(42). Our results and those of Chan *et al.* clearly exhibit a lower correlation than observed in human, however, our comparisons are done at the ˜200 kb scale where there is more variation in recombination rate (see below, **Table 3**). Furthermore, empirical-empirical correlations between different populations of *D. pseudoobscura* are only moderately higher than the LD-based-empirical correlations (**Table 2**), suggesting that the LD-based maps are capturing much of the recombination variation in these species.

**Table 2.** Empirical recombination rate comparisons. Empirical recombination rates were obtained from McGaugh *et al* 2012. Intervals were condensed between each pairing to correctly assess recombination rates as per McGaugh *et al* 2012.

|  | **Interval size**<br>**Average,**<br>**Median (Mb)** | **Number of**<br>**intervals** | **Correlation**<br>**(Pearson's r)** |
| --- | --- | --- | --- |
| **2** |  |  |  |
| *D. pseudoobscura* Flagstaff – Pikes Peak | 0.262, 0.170 | 115 | 0.718<br>p<0.0001 |
| *D. pseudoobscura* Flagstaff - *D. miranda* | 0.247, 0.152 | 115 | 0.621<br>p<0.0001 |
| *D. pseudoobscura* Pikes Peak- *D. miranda* | 0.235, 0.150 | 123 | 0.669<br>p<0.0001 |
| **XRgroup6** |  |  |  |
| *D. pseudoobscura* Flagstaff – Pikes Peak | 0.232, 0.202 | 47 | 0.916<br>p<0.0001 |
| *D. pseudoobscura* Flagstaff - *D. miranda* | 0.228, 0.194 | 48 | 0.656<br>p<0.0001 |
| *D. pseudoobscura* Pikes Peak- *D. miranda* | 0.197, 0.164 | 60 | 0.617<br>p<0.0001 |
| **XRgroup8** |  |  |  |
| *D. pseudoobscura* Flagstaff – Pikes Peak | 0.388, 0.280 | 21 | 0.873<br>p<0.0001 |
| *D. pseudoobscura* Flagstaff - *D. miranda* | 0.297, 0.257 | 20 | 0.776<br>p<0.0001 |
| *D. pseudoobscura* Pikes Peak- *D. miranda* | 0.296, 0.249 | 23 | 0.612<br>p<0.0001 |

**Table 3.** LD-based comparison of recombination rates between *D. pseudoobscura* and *D. miranda.* The Pearson correlation coefficient and p-value for LD-based recombination rates of *D. pseudoobscura* and *D. miranda*. Recombination rates are averaged over a given interval size, first for the whole chromosome (all chromosome groups concatenated), then for each chromosome group individually. Dashes indicate not enough data to assess the correlation.

| Chromosome | Size (bp) | Scale |  |  |  |
| --- | --- | --- | --- | --- | --- |
|  |  | 50kb | 100kb | 250kb | 500kb |
| **2** | 30794191 | 0.479<br>p<0.0001 | 0.499<br>p<0.0001 | 0.535<br>p<0.0001 | 0.510<br>p<0.0001 |
| **4** |  | 0.617<br>p<0.0001 | 0.668<br>p<0.0001 | 0.710<br>p<0.0001 | 0.786<br>p<0.0001 |
| **group1** | 5278962 | 0.365<br>p<0.0001 | 0.409<br>p=0.0024 | 0.403<br>p=0.0697 | - |
| **group2** | 1235211 | 0.424<br>p=0.0347 | - | - | - |
| **group3** | 11692073 | 0.660<br>p<0.0001 | 0.720<br>p<0.0001 | 0.774<br>p<0.0001 | 0.842<br>p<0.0001 |
| **group4** | 6587037 | 0.593<br>p<0.0001 | 0.643<br>p<0.0001 | 0.762<br>p<0.0001 | - |
| **XL** |  | 0.462<br>p<0.0001 | 0.485<br>p<0.0001 | 0.557<br>p<0.0001 | 0.669<br>p<0.0001 |
| **group1a** | 9151740 | 0.592<br>p<0.0001 | 0.608<br>p<0.0001 | 0.702<br>p<0.0001 | 0.930<br>p<0.0001 |
| **group1e** | 12523135 | 0.168<br>p=0.0077 | 0.182<br>p=0.0418 | 0.173<br>p=0.2303 | 0.151<br>p=0.4705 |
| **group3a** | 2690836 | 0.730<br>p<0.0001 | 0.790<br>p<0.0001 | 0.822<br>p=0.0019 | - |
| **group3b** | 386183 | - | - | - | - |
| **XR** |  | 0.529<br>p<0.0001 | 0.617<br>p<0.0001 | 0.654<br>p<0.0001 | 0.652<br>p<0.0001 |
| **group3a** | 1468912 | 0.557<br>p=0.0014 | 0.673<br>p=0.0059 | - | - |
| **group5** | 740492 | 0.039<br>p=0.8898 | - | - | - |
| **group6** | 13314419 | 0.532<br>p<0.0001 | 0.638<br>p<0.0001 | 0.693<br>p<0.0001 | 0.717<br>p<0.0001 |
| **group8** | 9212921 | 0.614<br>p<0.0001 | 0.720<br>p<0.0001 | 0.756<br>p<0.0001 | 0.716<br>p<0.0001 |

## Recombination rates between species

### Broad scale recombination rate patterns

Present-day empirically assayed recombination rate was previously shown to be conserved at broad scales (˜200kb) between *D. pseudoobscura* and *D. miranda* (**Table 2**) (63), and with our LD-based recombination maps, we recapitulate and extend these results. We detect and confirm genome wide broad scale conservation of recombination rates between species (**Table 3**, **Fig. 2, S2-S4 Figs**), but with several notable exceptions. First, *D. miranda* rates remain elevated relative to *D. pseudoobscura*, supporting the conclusion that the putative global recombination modifier identified in empirical work persists species-wide in *D. miranda* (63). Estimates of the effect of the modifier are quite similar between empirical and LD-based data, recombination rate is 1.26x higher on chromosome 2 in *D. miranda* (compared to 1.28-1.32 higher from empirical work), and 1.71x higher on chromosome XR (empirical: 1.4-1.47). Moreover, this difference persists despite a difference in effective population sizes: *D. pseudoobscura* N_e_ is thought to be several times that of *D. miranda* (70-72).

**Fig. 2:**
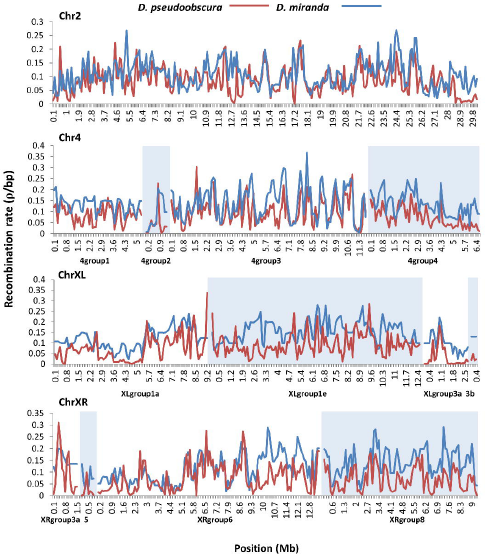
A comparison of LD-based estimations of recombination rate between two species. LD-based *D. pseudoobscura* recombination rates (red) are plotted with LD-based *D. miranda* recombination rates (blue) across the genome. Average rates were calculated for 100kb window sizes. Chromosomes 4, XL, and XR are split in to multiple groups, each labeled on the x-axis, according to the reference assembly for *D. pseudoobscura* (denoted with alternating background color). All recombination rates are reported in ρ/bp. Error bars are not shown for clarity.

Another pattern apparent in the broad scale recombination landscape is that some chromosome segments exhibit a weaker correlation of recombination rate between species, potentially indicative of regional differences. For example, chromosome arm XL group1e is a clear outlier in the comparison between *D. pseudoobscura* and *D. miranda* (**Table 3, Fig. 2, S2-S4 Figs**). Interestingly, the species differ by two inversions on the XL chromosome arm: a large 12 Mb inversion that corresponds almost precisely with group1e and a small, ˜4Mb inversion that took place following the large inversion at the distal end of group1e (73, 74). This is suggestive that an inversion caused historic broad scale changes in recombination rate.

### Fine scale recombination rate patterns

While recombination rates are generally conserved at broad scales, they are divergent at fine scales in the few organisms in which they have been examined. Analysis of fine scale recombination in Drosophila has been limited to isolated genomic regions in *D. melanogaster* (56-58, 75) and *D. pseudoobscura* (55, 63, 76), and has not been compared between species. With LD-based maps, we are able to characterize the genome-wide fine-scale recombination landscape in *D. pseudoobscura* and *D. miranda* for the first time, and to test whether turnover in fine scale recombination rates occurs outside of mammals.

By comparing recombination maps across different scales, variation and amplitude in recombination rate appears to increase at finer scales: for every chromosome assembly but one, the correlation coefficient between the species was weaker in 50kb windows than in every larger window size (100kb, 250kb, or 500kb: see **Table 3**). This phenomenon was previously noted in several Drosophila recombination studies (58, 63). In McGaugh *et al.* (63), the 20kb recombination map revealed recombination rates ranging from 3.5–21.2 cM/Mb, while the ˜200kb map measured rates of 5.6 and 4.4 cM/Mb covering the same region(17.5–17.7 Mb of chromosome 2). If we examine this same region in the LD-based maps (**S7 Fig.**), at position 17.5 Mb with resolution of 50kb, the recombination rate in *D. pseudoobscura* is 0.32 ρ/bp, while in the 500kb resolution map, the recombination rate is 0.16 ρ/bp, mirroring the empirical results. More generally, recombination rates peak at 0.32 ρ/bp in the 50kb chromosome 2 map for both species, and top out at 0.20 ρ/bp (*D. miranda*) and 0.14 ρ/bp (*D. pseudoobscura*) in the 1 Mb chromosome 2 map. It follows that correlations of recombination rate between two species are significantly higher at 500kb scale than at 50 kb scale (p=0.0094), illustrating stronger conservation at broad scales (**Table 3**, **S2-S4 Figs**). It appears that even in the absence of *Prdm9* and hotspots, there is less constraint at finer scales, although the mechanism behind this in Drosophila remains unclear.

While our study achieves the finest scale genome wide recombination map of both *D. pseduuobscura* and *D. miranda* to date, our resolution is limited biologically due to a lower SNP density in *D. miranda*, unfortunately restricting the level at which we can compare rates between species. The effective population size of *D. miranda* is estimated to be up to five times smaller than *D. pseudoobscura*, and number of segregating sites almost half of *D. pseudoobscura* (θ_pse_ = 0.01, θ_mir_=0.0058). Because statistical programs like LDhelmet depend on SNP density to estimate recombination, we are prohibited from examining recombination rates at a scale finer than 50kb in *D. miranda*.

However, we were still able to examine recombination rates at finer scales in *D. pseudoobscura*, and identified several putative hotspots in a similar manner to Chan *et al.* (64). Chan *et al.* detected 21 total hotspots in two different populations of *D. melanogaster* and were able to independently verify 10. In *D. pseudoobscura*, we discovered 19 regions between 500-5000bp where the recombination rate was more than ten times greater than the chromosomal recombination average (**S10 Fig.**). However, we did not verify these *D. pseudoobscura* hotspots independently with alternative means, so their presence should be seen as provisional, and these isolated hotspots remain both far less in total number and in magnitude than those seen in other organisms. This conclusion remains consistent with previous data, confirming that the recombination landscape of Drosophila possesses fine scale variation, but does not exhibit recombination hotspots like those found in humans, yeast, and others.

### The distribution of recombination across the genome

Looking at the proportion of recombination that occurs in a proportion of sequence reveals the degree to which recombination events cluster across the genome. In humans, recombination is highly punctate, with about 80% of recombination happening in less than 20% of the sequence (77), or a Gini coefficient of approximately 0.8 (59, 78). As previously discussed, Drosophila lack hotspots characteristic of other organisms (55, 57, 58), although fine scale variation has been identified (see above), and Chan *et al.* even detected a handful of more typical hotspots (64). This is reflected in *D. pseudoobscura*’s intermediate Gini coefficient of approximately 0.50, or about 80% of recombination occurring in 50% of the sequence (**Fig. 3**). This is similar to the Gini coeffiecient estimated for *D. melanoagaster* (0.47), falling between *C. elegans* (0.28) and *S. cerevisiae* (0.64) (59). This supports the hypothesis that recombination in Drosophila is more evenly distributed across the genome, with no region experiencing a near-complete absence of recombination, as compared to humans which exhibit large stretches of essentially no recombination events (77).

**Figure 3:**
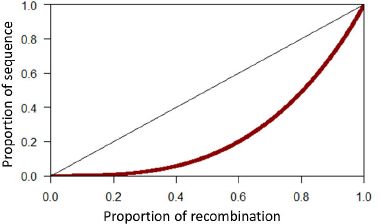
The concentration of fine-scale recombination rate in *D. pseudoobscura*.

Another way to examine the distribution of recombination is to examine recombination rate near genomic features, such as transcription start sites (TSS). As previously discussed, organisms without *Prdm9* exhibit an increase in recombination around TSS, most likely due to reduced nucleosome occupancy in these regions. However, in *D. pseudoobscura*, recombination is reduced around TSS and adjacent regions up to 35 kb away (**Fig. 4**). This is supported by findings in *D. melanogaster* as well (64), although the reduction in rates appears to persist for greater distances in *D. pseudoobscura*. Similarly, we see a slight reduction in recombination rate at the 5’ end of genes, which marginally increases with distance from the start of the coding region (**S11 Fig.**). Recombination rate within a gene (0.060 ρ/bp) is notably decreased from the average recombination rate of 0.098 ρ/bp.

**Figure 4:**
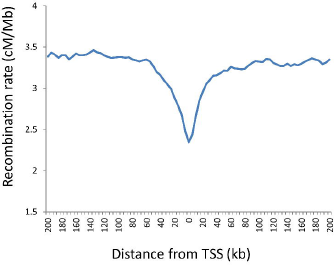
Recombination near transcription start sites in *D. pseudoobscura*. Recombination rate was averaged for 5kb windows for 200kb flanking transcription start sites (TSS) across the genome.

### Patterns of linked selection

One of the most ubiquitous patterns in molecular evolution is the association between recombination rate and nucleotide diversity, first observed in *Drosophila melanogaster* and since reproduced in many species, including several additional species of Drosophila (6, 79). The association with diversity, but not nucleotide divergence, has been interpreted as evidence that selective sweeps and/or background selection are eliminating diversity in regions of low recombination (80-85). Previous research has observed these associations at scales ranging from 50kb to 1.5 Mb, but failed to detect an association below 50kb (18, 58, 79, 86-88). This could potentially be explained by the size of the footprint of a selective event, such as a selective sweep, which may be undetectable at fine scales. To characterize the correlation between recombination and diversity at varying scales, we assessed diversity at four-fold degenerate positions binned into windows ranging from 5kb to 1Mb (**S12 Fig.**). Similar to Singh *et al.* (58), we detect a weak correlation between recombination and diversity at very fine scales, but this correlation increases in strength with window size so that at 1 Mb, recombination explains 36% of the variance in diversity. This pattern can be explained in part by the smoothing of stochasticity in diversity and recombination rate measures in large windows, and is potentially confounded by the method of recombination estimation depending on SNP density, although this method has been used for this purpose many times (64, 83, 89). We were able to re-test the correlation for a sample of windows using thinned SNP data (excluding 50% of SNPs), and did not find it to alter our results. We therefore conclude that patterns of linked selection are indeed apparent in *D. pseudoobscura*, and the power of this selection is strongest at large scales.

## Conclusions

With our genome wide fine scale recombination maps for two closely related species, we were able to identify both shared and exclusive patterns of recombination across different taxa. For example, without hotspots, it was unclear whether Drosophila would recapitulate the pattern observed in several other species, in which broad scale recombination rates are more conserved than fine scale recombination rates. While we could not compare rates at the 1-2kb level, by examining correlations at levels from 500kb to 50kb, we can conclude that recombination rates are less conserved at fine scales across the genome, one of the only studies to show this phenomenon exists outside of mouse and human. This raises interesting implications about selection pressures on recombination that persist across distantly related taxa with widely different recombination landscapes, supporting a relaxed constraint on rates at finer scales.

To illuminate potential strain variation or local changes in recombination rate over time, we can compare historical and present day recombination rates between species (**Tables 1**-**3**). Present recombination rates between species are moderately well correlated (**Table 3**), suggesting that recombination rate has not experienced drastic changes since these species diverged, although this analysis is limited to chromosome 2 and parts of XRgroup6 and XRgroup8. Comparing historical rates (LD-based) between species at a similar scale (**Table 2**), we see a decrease in correlation strength compared to present day, some regions quite dramatically. Generally, this likely represents variability across strains, as the LD-based recombination rates portray a population average. In specific regions like 4group1 and XLgroup1e, which seem to exhibit very different historical recombination patterns between species, this may indicate species specific local changes to recombination rate. As discussed above, one potential driver of this change is chromosomal rearrangements, such as inversions. Fittingly, an inversion between *D. miranda* and *D. pseudoobscura* overlaps almost exactly with the region of greatest diverge on chromosome arm XLgroup1e. This phenomenon has been previously documented in human-chimpanzee broad scale correlations, where inverted regions showed a lower correlation in recombination rate than non-inverted regions (21), supporting the conclusion that inversions influence broad scale patterns of recombination across multiple species.

Another potential modifier of recombination, originally implicated in empirical work (63), was confirmed here by observing genome wide elevated recombination in historical recombination rates of *D. miranda* despite a much lower effective population size. This new data provides compelling evidence that the predicted recombination modifier is a species wide trait. Of course, modifiers of recombination are well known to Drosophila (90, 91), and the theory of conditions that favor increased (or decreased) recombination is rich (92-99). The selection pressures that *D. miranda* has faced are unknown, and unfortunately, determining the locus or loci responsible for an increase in recombination in this species is not possible through traditional mapping methods, as *D. pseudoobscura* and *D. miranda* do not interbreed.

Of course one of the most recent, well-studied determinants of recombination, *Prdm9*, is absent in Drosophila, just one of many other differences that set Drosophila recombination apart from other organisms. As has already been discussed, Drosophila lack hotspots of recombination, and exhibit a more consistent distribution of recombination than yeast, mouse, and human. Additionally, recombination is decreased around the start of genes, in contrast to other organisms without *Prdm9*. These results add further evidence to long known differences in Drosophila meiosis, which can largely be attributed to the loss of recombination in males. Whether there’s a link between the evolution of sex-specific recombination and the patterns observed more recently is unclear, but the decreased cost of sequencing and computational approaches like LDhelmet may help elucidate this question by characterizing the recombination landscape of other sex-specific recombination systems, among other questions such as why sex-specific recombination has evolved so many times, and why recombination hotspots were lost in the lineage leading to Drosophila when they persist across fungi, plants, and metazoans.

In conclusion, the creation of recombination maps will continue to illuminate new perspectives on the evolution of recombination, shedding light on novel determinants of recombination, patterns of selection, and genome evolution.

## Materials and Methods

### Population sequencing and variant calling

We used 11 whole genome sequences of *D. pseudoobscura* (MV2-25 (reference), Mather32, MSH24, MSH9, TL, PP1134, PP1137, AFC12, FLG14, FLG16, FLG18) previously sequenced and analyzed by McGaugh et al. (63) and which are available at http://pseudobase.biology.duke.edu and GenBank (accession numbers: AFC12 SRX091462, Flagstaff14 SRX091308, Flagstaff16 SRX091303, Flagstaff18 SRX091310, Mather32 SRX091461, MatherTL SRX091324, MSH9 SRX091465, MSH24 SRX091463, PP1134 SRX091323, PP1137 SRX091311). Additionally, we used 11 whole genome sequences of *D. miranda* (Lines: MA28, MAO101.4, MAO3.3, MAO3.4, MAO3.5, MAO3.6, ML14, ML16, ML6f, SP138, and SP235), previously sequenced and provided by Doris Bachtrog’s lab. There is little to no population structure present in *D. pseudoobscura*, allowing us to treat flies collected from various regions across the North American west as panmictic (100-102). All sequences of *D. pseudoobcura* and *D. miranda* were aligned to the *D. pseudoobscura* reference sequence v2.9 using bwa (103). Variants were called for the *D. miranda* genome sequences using the Genome Analysis Tool Kit (GATK) v2.7-2 to remove duplicates and locally realign reads (104, 105), and samtools v0.1.18 to call single nucleotide variants (106). We used custom Perl scripts to convert the variant output files into aligned species-specific individual chromosome FASTA files for input to LDhelmet, see below (2, 4group1, 4group2, 4group3, 4group4, XLgroup1a, XLgroup1e, XLgroup3a, XLgroup3b, XRgroup3a, XRgroup5, XRgroup6, XRgroup8). Note that chromosome 3 was not included in the analysis due to known segregating inversions in *D. pseudoobscura* (107), and chromosome 4group5 was excluded due to poor sequence coverage.

### Recombination estimates

Empirical estimates of recombination for *D. pseudoobscura* and *D. miranda* were obtained from McGaugh et al. (63)(**Table 2**). Population sequencing based estimates of recombination were determined using the LDhelmet program (64), a statistical approach designed for Drosophila which estimates the population recombination parameter ρ using a reversible jump Markov Chain Monte Carlo mechanism (rjMCMC), as in LDhat (3, 4). The program calculates ρ=4Ner (where Ne is the effective population size and r is the recombination rate per generation) to estimate the amount of recombination needed in the population to produce the observed levels of linkage disequilibrium under a given model. LDhelmet is specifically tailored to address issues in Drosophila that may be problematic for programs such as LDhat, like a magnitude fold higher background recombination rate, higher SNP density, and a large portion of the genome influenced by positive selection (82, 108). The program was run individually for each chromosome of each species using default parameters, with the exception that we estimated theta from our dataset (θ_pse_ = 0.01, θ_mir_=0.0058) and created our own mutation transition matrices. Following the procedure of Chan et al., we ran the program for 1,000,000 iterations after a burn-in of 100,000 iterations, and used a conservative block penalty of 50.

To compare empirical recombination estimates to LD-based recombination estimates, the recombination estimate from LDhelmet was corrected for distance and averaged over a given interval. The empirical chromosomal recombination average (cM/Mb) was divided by the total average LD-based recombination rate for a chromosome to get a conversion factor. Each interval’s average LD-based estimate was multiplied by the conversion factor to get an approximation of recombination rates in the units centiMorgan per Megabase (cM/Mb) per Chan *et. al* (64).

### Analyses of genomic correlates

#### Transcription start sites (TSS)

Locations of *D. pseudoobscura* transcription start sites (TSS) were obtained from Main *et al.* (109). The recombination estimates from LDhelmet were averaged over 5000bp intervals for the 200kb regions flanking either side of a TSS. Genome wide data was aggregated to create **Figure 3**. We confirmed that the effect seen around TSS is not due to a lower SNP density caused by conservation of promoter regions, by comparing SNP density in 20 kb regions flanking TSS to 20 kb regions where no genes are present.

#### Proportion recombination

A Perl script was used to sort recombination rates and distances to build a list of the proportion of recombination occurring in the proportion of sequence (Laurie Stevison, unpublished). This data was plotted using a Lorenz curve from the R package “ineq” (Achim Zeileis, Christian Kleiber). The Gini coefficient was also calculated using this package as in Kaur *et al.* (59).

#### Introns and Exons

Locations of exons and introns and relative positions in a gene were extracted from *D. pseudoobscura* v2.9 annotations from FlyBase (110). The recombination estimate from LDhelmet was corrected for distance and averaged over the given interval, then aggregated to give genome wide totals for each exon and intron position within a gene.

#### Nucleotide diversity

We calculated pairwise nucleotide diversity (π) at four-fold degenerate sites using custom Perl scripts, excluding sites where an insertion or deletion was found in any line.

#### Data archiving

All datasets used to create the figures or tables in this study are archived in Dryad, doi:10.5061/dryad.sh5c3.

## Acknowledgements

Thanks To L. Stevison for helpful comments on the preparation of this manuscript. We are grateful to D. Bachtrog for sharing the *D. miranda* whole genome sequences with us. Our sincere thanks to Andrew H. Chan for his generous guidance and assistance with LDhelmet throughout the project.

## Supporting Information

**S1 Fig.**
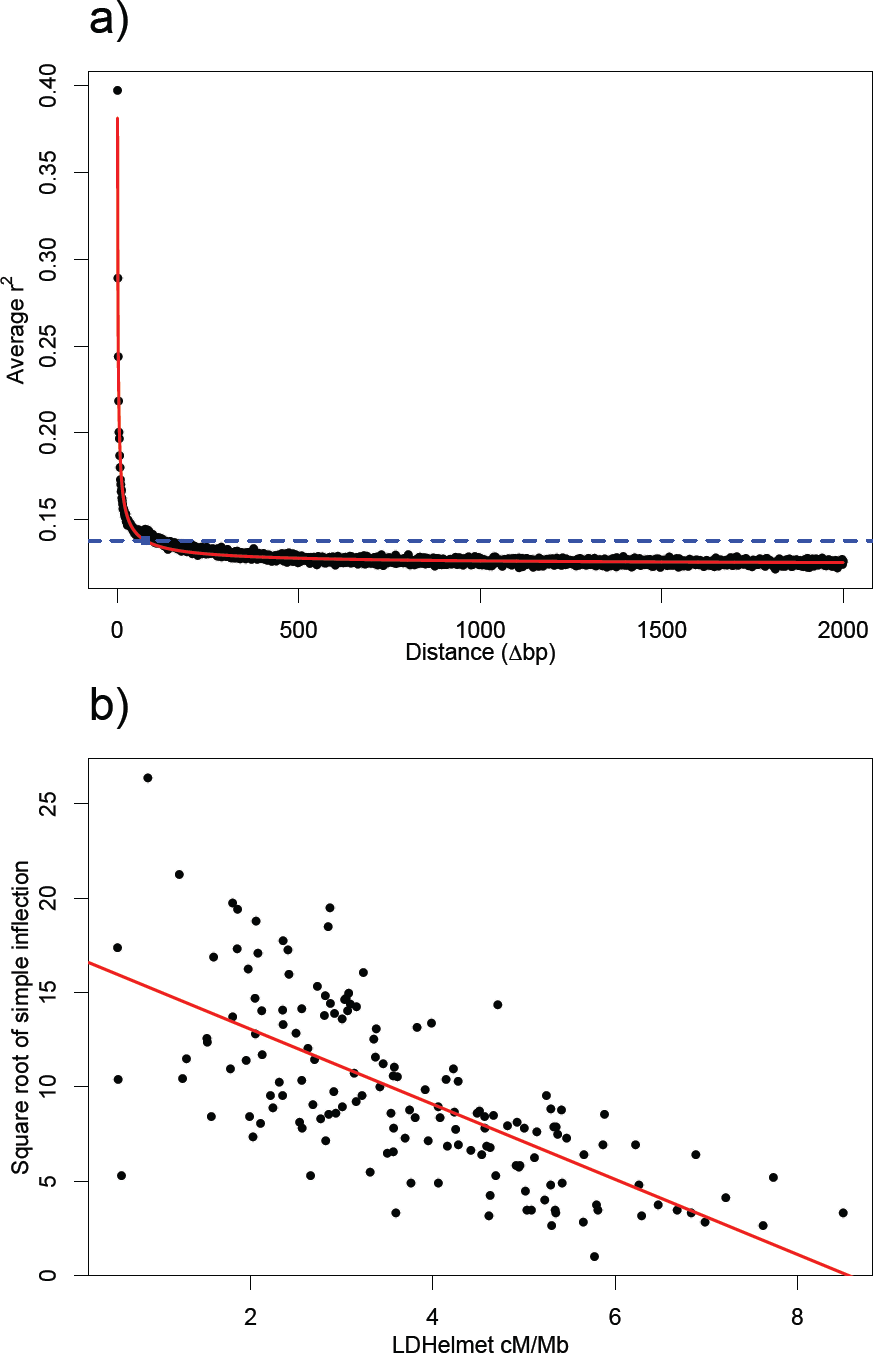
A comparison of the LDhelmet estimate of recombination rate with an alternative simple direct measure of linkage disequilibrium decay. a) Average r^2^ as a function of distance between SNPs was generated for all polymorphic base pairs on chromosome 2. The best-fit line (red) was generated using a five-parameter log-logistic function. LD was inferred to have decayed when the best-fit line came to a value 10% higher than background r^2^ (blue line, calculated as the average r2 of SNPs at distance 1500 – 2000bp from each other), b) Correlation between linkage disequilibrium calculated through LDhelmet and the square root of the nucleotide distance at which LD decayed as estimated through the method described in a).

**S2 Fig.**
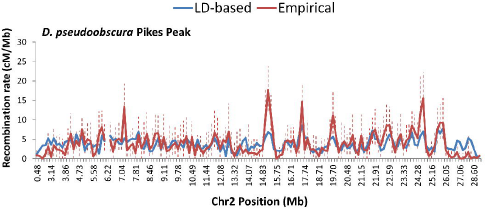
A comparison of empirical and LD-based estimations of recombination rate for *D. pseudoobscura* Pikes Peak, Chr2. Empirical recombination rates (red) are plotted with LDhelmet derived recombination rates (blue) across chromosome 2 for *D. pseudoobscura* Pikes Peak. All recombination rates are reported in cM/Mb. Error bars depict 95% Confidence Intervals for the empirical (red, dashed), and LD-based (black, solid).

**S3 Fig.**
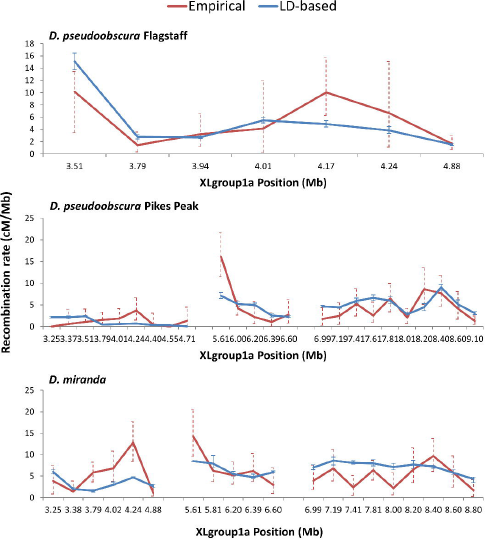
A comparison of empirical and LD-based estimations of recombination rate for ChrXL. Empirical recombination rates (red) are plotted with LDhelmet derived recombination rates (blue) across portions of chromosome XL group1a for *D. pseudoobscura* Flagstaff (top), *D. pseudoobscura* Pikes Peak (middle), and *D. miranda* (bottom). All recombination rates are reported in cM/Mb. Error bars depict 95% Confidence Intervals for the empirical (red, dashed), and LD-based (blue, solid).

**S4 Fig.**
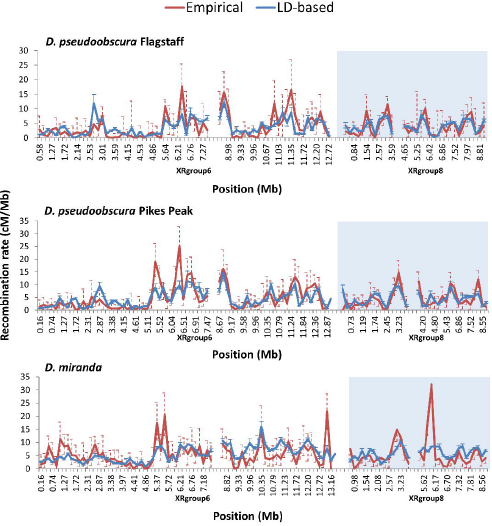
A comparison of empirical and LD-based estimations of recombination rate for ChrXR. Empirical recombination rates (red) are plotted with LDhelmet derived recombination rates (blue) across portions of chromosome XR group 6 and 8 for *D. pseudoobscura* Flagstaff (top), *D. pseudoobscura* Pikes Peak (middle), and *D. miranda* (bottom). All recombination rates are reported in cM/Mb. Error bars depict 95% Confidence Intervals for the empirical (red, dashed), and LD-based (blue, solid).

**S5 Fig.**
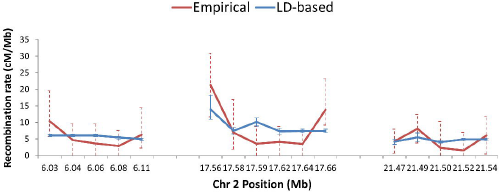
A comparison of empirical and LD-based estimations of fine scale recombination rate. Empirical recombination rates (red) are plotted with LDhelmet derived recombination rates (blue) for three regions of 20kb intervals on chromosome 2 of *D. pseudoobscura* Flagstaff. All recombination rates are reported in cM/Mb. Error bars depict 95% Confidence Intervals for the empirical (red, dashed), and LD-based (blue, solid).

**S6 Fig.**
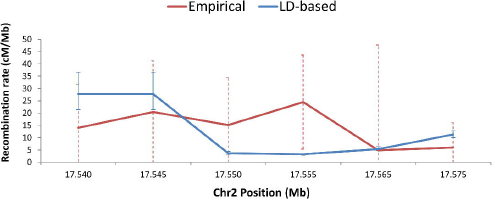
A comparison of empirical and LD-based estimations of super-fine scale recombination rate. Empirical recombination rates (red) are plotted with LDhelmet derived recombination rates (blue) for one region of 5kb intervals on chromosome 2 of *D. pseudoobscura* Flagstaff. All recombination rates are reported in cM/Mb. Error bars depict 95% Confidence Intervals for the empirical (red, dashed), and LD-based (blue, solid).

**S7 Fig.**
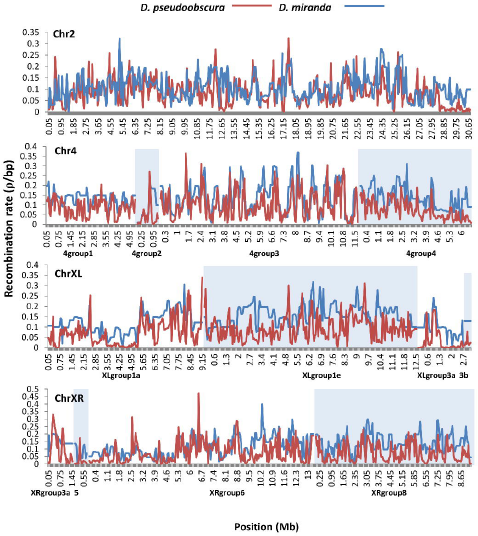
A comparison of LD-based estimations of recombination rate between two species at 50kb. LD-based *D. pseudoobscura* recombination rates (red) are plotted with LD-based *D. miranda* recombination rates (blue) across the genome. Average rates were calculated for 50kb window sizes. Chromosomes 4, XL, and XR are split in to multiple groups, each labeled on the x-axis, according to the reference assembly for *D. pseudoobscura* (denoted with alternating background color). All recombination rates are reported in ρ/bp. Error bars are not shown for clarity.

**S8 Fig.**
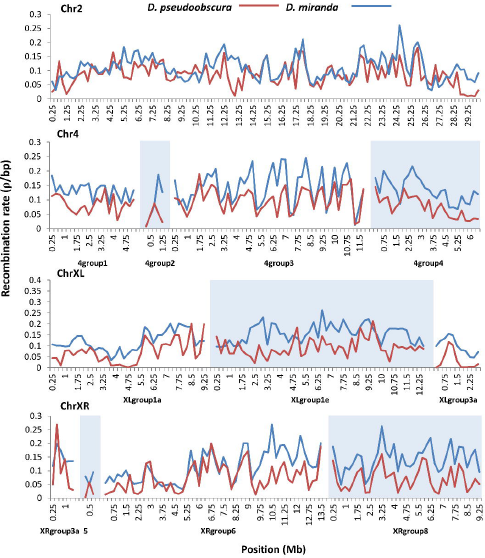
A comparison of LD-based estimations of recombination rate between two species at 250kb. LD-based *D. pseudoobscura* recombination rates (red) are plotted with LD-based *D. miranda* recombination rates (blue) across the genome. Average rates were calculated for 250kb window sizes. Chromosomes 4, XL, and XR are split in to multiple groups, each labeled on the x-axis, according to the reference assembly for *D. pseudoobscura* (denoted with alternating background color). Smaller chromosome groups are excluded due to lack of data. All recombination rates are reported in ρ/bp. Error bars are not shown for clarity.

**S9 Fig.**
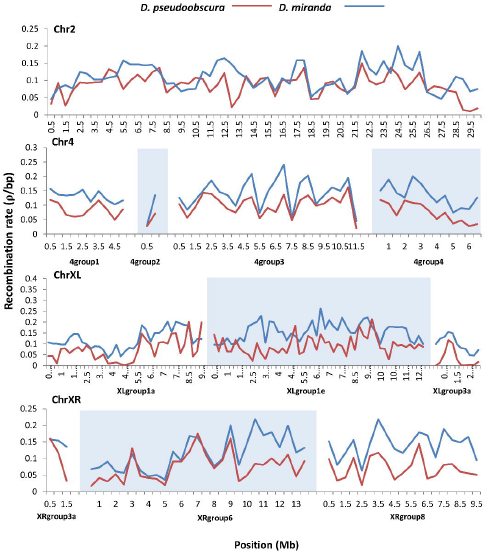
A comparison of LD-based estimations of recombination rate between two species at 500kb. LD-based *D. pseudoobscura* recombination rates (red) are plotted with LD-based *D. miranda* recombination rates (blue) across the genome. Average rates were calculated for 500kb window sizes. Chromosomes 4, XL, and XR are split in to multiple groups, each labeled on the x-axis, according to the reference assembly for *D. pseudoobscura* (denoted with alternating background color). Smaller chromosome groups are excluded due to lack of data. All recombination rates are reported in ρ/bp. Error bars are not shown for clarity.

**S10 Fig.**
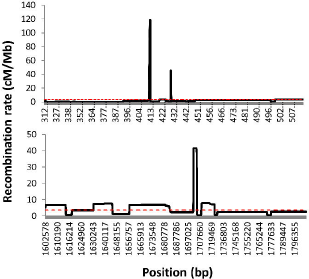
Recombination hotspots in *D. pseudoobscura*. Two recombination “hotspots” from chromosome 2 of *D. pseudoobscura* plotted with the background recombination rate (red, dashed). The region of increased recombination is shown with 100kb flanking sequence. Recombination rates are reported in cM/Mb.

**S11 Fig.**
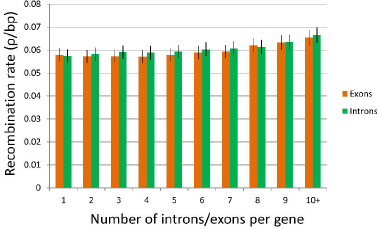
Recombination rate within genes of *D. pseudoobscura*. Recombination rates are slightly decreased at the start of genes. Genes were characterized by exon and intron number, then recombination rate was averaged over the genomic position of the exon or intron. Recombination rates were aggregated to give genome wide totals for each exon (orange) and intron(green) position within a gene. Recombination rates are reported in ρ/bp.

**S12 Fig.**
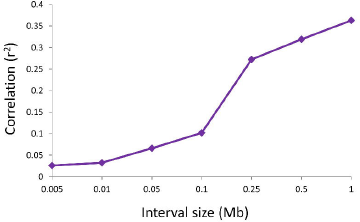
Linked selection over varying physical scales in *D. pseudoobscura*. Recombination rate was plotted with pairwise nucleotide diversity at four-fold degenerate positions across the genome of *D. pseudoobscura* with window sizes of 5kb, 10kb, 50kb, 100kb, 250kb, 500kb, and 1Mb. The coefficient of determination (r^2^) for each of these associations is plotted (5kb=0.0262, 10kb=0.0326, 50kb=0.0664, 100kb=0.1019, 250kb=0.2721, 500kb=0.3192, and 1Mb=0.3628), showing recombination increasingly explains the variance in nucleotide diversity at large physical scales.

